# Trogocytosis by *Entamoeba histolytica* mediates acquisition and display of human cell membrane proteins and evasion of lysis by human serum

**DOI:** 10.1101/487314

**Authors:** Hannah W. Miller, Rene L. Suleiman, Katherine S. Ralston

## Abstract

*Entamoeba histolytica* is the protozoan parasite responsible for amoebiasis. We previously showed that *E. histolytica* kills human cells through a mechanism that we termed trogocytosis (*trogo-:* nibble), due to its resemblance to trogocytosis in other organisms. In parasites, trogocytosis is used to kill host cells. In multicellular organisms, trogocytosis is used for cell-cell interactions in the immune system, in the central nervous system, and during development. Thus, nibbling is an emerging theme in cell-cell interactions both within and between species, and it is relevant to host-pathogen interactions in many different contexts. When trogocytosis occurs between mammalian immune cells, cell membrane proteins from the nibbled cell can be acquired and displayed by the recipient cell. In this study, we tested the hypothesis that through trogocytosis of human cells, amoebae acquire and display human cell membrane proteins. Here we demonstrate for the first time that through trogocytosis, *E. histolytica* acquires and displays human cell membrane proteins and that this leads to protection from lysis by human serum. Protection from human serum only occurs after amoebae have undergone trogocytosis of live cells, but not phagocytosis of dead cells. Likewise, mutant amoebae that exhibit a phagocytosis defect, but are unaltered in their capacity to perform trogocytosis, are nevertheless protected from human serum. Our studies are the first to demonstrate that amoebae can display human cell membrane proteins and suggest that acquisition and display of membrane proteins is a general feature of trogocytosis that is not restricted to trogocytosis between mammalian immune cells. These studies have major implications for interactions between *E. histolytica* and the immune system and also reveal a novel strategy for immune evasion by a pathogen. Since other microbial eukaryotes use trogocytosis for cell killing, our findings may apply to the pathogenesis of other infections.

**Author Summary:** *Entamoeba histolytica* is an intestinal parasite that causes amoebiasis, a potentially fatal diarrheal disease. Abscesses in organs outside of the intestine, such as the liver, can occur when amoebae are able to breach the intestinal wall and travel through the blood stream to other areas of the body. We previously showed that *E. histolytica* kills human cells by taking “bites” of human cell material in a process that we named trogocytosis (*trogo-:* nibble). Mammalian immune cells use trogocytosis to acquire proteins from other cells which impacts cell-cell communication. Here we tested the hypothesis that trogocytosis allows *E. histolytica* to acquire and display proteins from human cells allowing amoebae to survive in the blood stream. We demonstrate for the first time that through trogocytosis, *E. histolytica* acquires and displays human cell membrane proteins. We also demonstrate that trogocytosis of human cells allows amoebae to survive in human serum. These studies reveal a novel strategy for immune evasion by a pathogen and may apply to the pathogenesis of other infections.

## Introduction

*Entamoeba histolytica* is the protozoan parasite responsible for amoebiasis, a potentially fatal diarrheal disease. Amoebiasis occurs worldwide and is most prevalent in developing countries, in areas with poor sanitation and high malnutrition [1–3]. For instance, a recent study found that nearly 80% of infants living in an urban slum in Bangladesh had been infected with *E. histolytica* by two years of age [4]. The infection has a wide range of clinical symptoms that include asymptomatic infection, diarrhea, bloody diarrhea, and fatal abscesses outside of the intestine. Bloody diarrhea arises when amoebic trophozoites (amoebae) invade and ulcerate the intestine. Amoebae that have invaded the intestine can then disseminate and cause abscesses in other tissues, most commonly in the liver. Although amoebic liver abscesses are rare, they are fatal if untreated. Little is known about the mechanisms that allow *E. histolytica* to evade immune detection and disseminate upon entering the bloodstream during invasive amoebiasis.

The parasite was named “histolytica” for its ability to damage tissue (*histo-:* tissue; *lytic-:* dissolving) [5–7]. Despite this name-giving property, how amoebae invade and damage tissues is not clear. The most well-known virulence factor is the amoeba surface D-galactose and N-acetyl-D-galactosamine (Gal/GalNAc) lectin [8,9], which mediates attachment to human cells and intestinal mucin [10–13]. Surface-localized and secreted cysteine proteases contribute to proteolysis of human substrates including mucin and extracellular matrix [10–13]. The profound cell killing activity of amoebae is likely to drive tissue damage. Amoebae can kill almost any type of human cell within minutes and direct contact with human cells is required for killing to occur [8,9]. Until recently, the accepted model was that the pore-forming amoebapores act as secreted toxins [14–17]. However, the contact-dependence of cell killing [8,9], and the lack of killing activity in cell lysates and supernatants [6,7,18], are not consistent with the presence of secreted toxins. Furthermore, transfer of amoebapores to human cells has not been demonstrated.

We previously established a new paradigm by showing that *E. histolytica* kills human cells through a mechanism that we termed trogocytosis (*trogo-:* nibble), due to its resemblance to trogocytosis in other organisms [19]. During trogocytosis, amoebae kill human cells by extracting and ingesting “bites” of human cell membrane and intracellular contents [19]. We defined that trogocytosis requires amoebic actin rearrangements [19]. It also requires signaling initiated by the Gal/GalNAc lectin, phosphatidylinositol 3-kinase (PI3K) signaling and an amoebic C2 domain-containing kinase (*EhC2PK*) [19]. By applying multiphoton imaging using explanted mouse intestinal tissue from fluorescent-membrane mice, we found that trogocytosis was required for tissue invasion, demonstrating relevance to pathogenesis [19].

Trogocytosis is not specific to *E. histolytica,* as it can be observed in other microbial eukaryotes as well as multicellular eukaryotes [20]. Examples in microbes include reports of trogocytosis by *Naegleria fowleri* [21] and by *Dictyostelium caveatum* [22]. In multicellular eukaryotes, trogocytosis is used for cell-cell communication and cell-cell remodeling. Trogocytosis plays roles in the immune system [23,24], in the central nervous system [25,26], and during development [27]. Additionally, intracellular bacteria exploit macrophage trogocytosis to spread from cell to cell [28]. It is not yet clear how trogocytosis can paradoxically be both a benign form of cell-cell interaction and a mechanism for cell-killing. The previous paradigm was that microbes engage trogocytosis for cell-killing, but trogocytosis in multicellular organisms was believed to be a benign form of cell-cell interaction. However, recent reports have now shown that neutrophils can use trogocytosis to kill parasites [29], and that neutrophils and macrophages can use trogocytosis to kill cancer cells in a form of antibody-dependent cell-mediated cytotoxicity [30,31]. Trogocytosis is therefore likely to be a conserved, fundamental form of eukaryotic cell-cell interaction that can be cytotoxic or benign, depending on the context.

One intriguing outcome of trogocytosis between mammalian immune cells is that it changes the makeup of cell surface proteins on both the donor and the recipient cell. The nibbling cell is able to display the acquired membrane proteins from the nibbled cell on its surface [24,32]. Acquired membrane proteins appear as foci or patches on the recipient cell. This allows the recipient cell to take on new properties that impact its subsequent interactions with other cells [24,32]. For instance, uninfected dendritic cells can acquire and display pre-loaded major histocompatibility complex class II (MHC II) molecules by nibbling infected dendritic cells, and thus they can present peptides from microbes they have not directly encountered, which has been termed “cross-dressing” [24]. Transferred molecules are not limited to MHC complexes as induced regulatory T cells can acquire cluster of differentiation (CD) molecules from mature dendritic cells including CD80 and CD86 [33]. It has also been shown that monocytes, NK cells, and granulocytes can acquire CD22, CD19, CD21, and CD79b from antibody-opsonized B cells [34]. In addition to allowing the nibbling cell to display newly acquired membrane proteins, trogocytosis also affects the makeup of surface proteins on the nibbled cell. Since membrane fragments are removed from the nibbled cell, trogocytosis affects the nibbled cell by effectively downregulating surface proteins [35].

Since mammalian immune cells acquire and display membrane proteins through trogocytosis, we hypothesized that amoebae may acquire and display human cell membrane proteins during trogocytosis. Amoebic display of human proteins would have significant implications for host-pathogen interactions. We predicted that one outcome of amoebic human cell protein display could be the inhibition of lysis by human complement. Previous studies have suggested that amoebae become more resistant to complement after interacting with host cells or tissues, and that complement resistance involves proteins on the amoeba surface. For example, amoebae became more resistant to complement after co-incubation with erythrocytes, and an antibody directed towards an erythrocyte membrane protein reacted with the amoeba surface after erythrocyte co-incubation [36]. Animal-passaged amoebae are more resistant to complement lysis than control amoebae [37], and treatment of complement-resistant amoebae with trypsin renders amoebae complement-sensitive [38].

Here we show that *E. histolytica* acquires and displays human cell membrane proteins. Acquisition and display of human cell membrane proteins requires actin and is associated with subsequent protection from lysis by human serum. Protection from human serum further requires actin and direct contact between amoebae and human cells. Protection from human serum occurs after amoebae have undergone trogocytosis, but not phagocytosis, suggesting protection is not generally associated with ingestion. Furthermore, mutant amoebae that are deficient in performing phagocytosis but not trogocytosis are still protected from human serum. Collectively, these findings support that amoebae acquire and display human cell membrane proteins through trogocytosis and that this leads to protection from lysis by human serum complement. These studies have major implications for interactions between *E. histolytica* and the immune system. Display of human cell proteins acquired during trogocytosis is a novel strategy for immune evasion by a pathogen. Since other microbial eukaryotes use trogocytosis for cell killing, this may apply to the pathogenesis of other infections.

## Results

### Amoebae acquire and display human cell membrane proteins

We first asked whether trogocytosis by *E. histolytica* could result in transfer of human cell membrane proteins to the cell membrane of the amoeba. Human Jurkat T cells were surface-biotinylated and then coincubated with amoebae. After co-incubation, cells were fixed and labeled with fluorescently-conjugated streptavidin (Fig 1A). Since cells were not permeabilized, this approach required human cell proteins to be surface-exposed and to retain correct orientation. After five minutes of co-incubation, patches of streptavidin-labeled human cell proteins were detected on the surface of amoebae (Fig 1B – 1C, arrows). Similar to immune cell “cross-dressing” [24], the biotin-streptavidin label appeared as patches or foci on the amoeba surface. To track an individual human cell membrane protein, immunofluorescence was used to detect human major histocompatibility complex class I (MHC I) (Fig 1D). Following co-incubation, cells were fixed without permeabilization, and MHC I was detected using a monoclonal antibody. Comparable to the biotin-streptavidin labeling experiments, MHC I was detected in patches on the surface of amoebae after five minutes of co-incubation (Fig 1E – 1F). Together, these results suggested that human cell membrane proteins were acquired and displayed by amoebae following co-incubation with live human cells.

**Fig 1.**
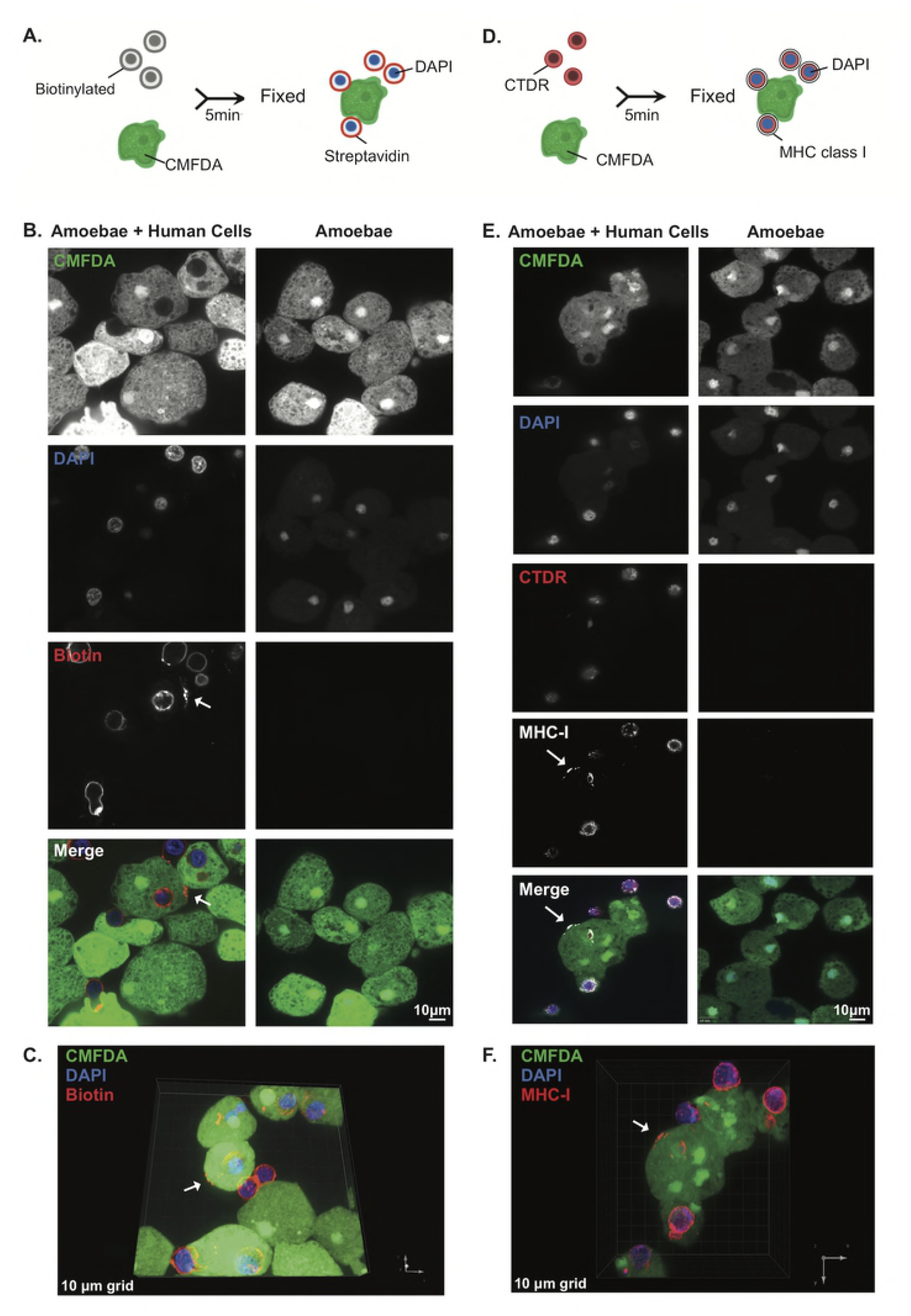
Following interaction with human cells, human cell membrane proteins are displayed by amoebae. **(A)** Human cell membrane proteins were labeled with biotin prior to co-incubation with CMFDA-labeled amoebae. Cells were co-incubated for 5 minutes and immediately fixed. Following fixation, samples were labeled with fluorescently-conjugated streptavidin and DAPI. **(B)** Representative images of amoebae incubated alone or co-incubated with biotinylated human cells. Amoebae are shown in green and streptavidin is shown in red. Nuclei are shown in blue. Arrow indicates a patch of biotin-streptavidin localized to the amoeba surface. **(C)** 3D rendering of Z stack images taken from panel B. Arrow indicates transferred biotin. **(D)** Human cells were labeled with cell tracker deep red (CTDR) prior to co-incubation with CMFDA-labeled amoebae. Cells were co-incubated for 5 minutes and immediately fixed. Following fixation, samples were labeled with DAPI and MHC-I was detected using immunofluorescence. **(E)** Representative images of amoebae incubated alone or co-incubated with CTDR-labeled human cells. Amoebae are shown in green, human cell cytoplasm is shown in red, MHC-I is shown in yellow and nuclei are shown in blue. Arrow indicates MHC-I present on the amoeba surface. **(F)** 3D rendering of Z stack images taken from E. Arrow indicates transferred MHC-I. For panels B-F, images were collected from 4 independent experiments.

### Acquisition and display of human cell membrane proteins requires actin

Trogocytosis by *E. histolytica* requires actin rearrangements and is inhibited by treatment with cytochalasin D [19]. Therefore, we next asked whether transfer of human cell membrane proteins required actin. Imaging flow cytometry was used to quantitatively analyze displayed biotinylated human cell membrane proteins on the amoeba surface. It was important to distinguish between amoebae that displayed human cell membrane proteins and amoebae that were attached to intact, extracellular human cells. While the latter may also display human cell membrane proteins, we focused our analysis on images that lacked extracellular human cells as this allowed for the highest stringency in quantifying displayed human cell membrane proteins. Since human cell nuclei are not internalized by amoebae during trogocytosis [19], human cell nuclei were fluorescently labeled, and this was used to gate images that contained or lacked extracellular human cells.

Human cell nuclei were labeled with Hoechst, and human cell membrane proteins were biotinylated prior to co-incubation with CellTracker Green CMFDA Dye (CMFDA)-labeled amoebae. After gating on single amoebae out of total cells (Fig 2A, S1 Fig), Hoechst staining was used to gate on images of amoebae with and without extracellular human cells (Fig 2B, D, F). Next, the extent of overlap of fluorescent-streptavidin and individual amoebae was quantified (Fig 2C, E). In the dimethyl sulfoxide (DMSO) treated control amoebae, 25% of amoebae contained patches of biotin labeling (Human Cell Nuclei-/Biotin+), while in the cytochalasin D treated amoebae, 5% of amoebae contained patches of biotin labeling (Fig 2E). From this, we concluded that amoebae acquire and display human cell membrane proteins through an actin-dependent process, consistent with trogocytosis.

### Interaction with human cells leads to protection from lysis by human serum

The acquisition and display of human cell membrane proteins has many potential implications for host-parasite interactions. One possible implication is in resistance to lysis by complement in human serum, particularly since it has been previously suggested that ingestion of human erythrocytes protects amoebae from lysis by human complement [36]. Amoebae preferentially perform trogocytosis on live human cells [19], therefore amoebae were incubated in the presence or absence of live human cells and then exposed to human serum (Fig 3A, S2 Fig). Using imaging flow cytometry, amoeba viability (Fig 3C – D), and trogocytosis were simultaneously measured (Fig 3B, S3 Fig). Amoebae that had interacted with live human cells, and had thus undergone trogocytosis and acquired human cell membrane proteins, were quantitatively protected from lysis by human serum (Fig 3C – 3D, S4 Fig). Among amoebae that had been incubated with human cells, amoebae that were lysed by human serum had generally undergone less trogocytosis than amoebae that survived exposure to human serum (Fig 3E). Therefore, trogocytosis is associated with subsequent protection from lysis by human serum.

**Fig 2.**
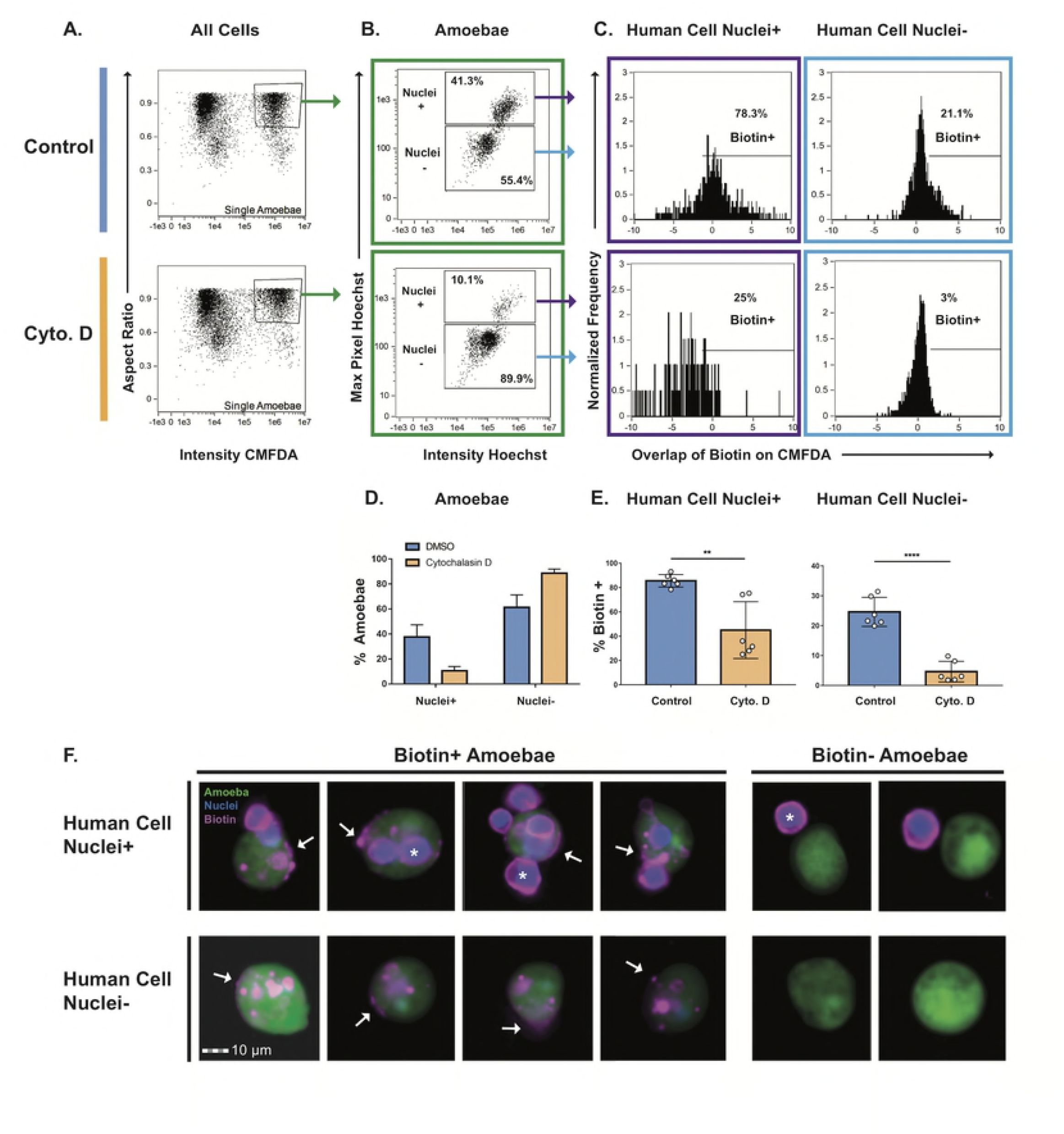
Acquisition of human cell membrane proteins is inhibited with cytochalasin D treatment. CMFDA-labeled amoebae were pre-treated with either cytochalasin D (Cyto. D) or DMSO (Control) and were then combined with Hoechst-labeled human cells. Immediately after co-incubation, cells were placed on ice to halt ingestion and stained with fluorescently-conjugated streptavidin. Samples were quantitatively analyzed using imaging flow cytometry, with 10,000 images collected for each sample. **(A)** Gate used to identify single amoebae from total cells. Focused cells were gated on single amoebae using aspect ratio and intensity of CMFDA fluorescence. **(B)** Representative plots of images with and without human cell nuclei (Hoechst high or low populations) are shown. The Hoechst high population contained images of amoebae with human cells and the Hoechst low population contained images of amoebae without human cells. **(C)** Overlap of biotin and CMFDA fluorescence was measured, and biotin positive images were gated. Representative plots of DMSO and cytochalasin D treated samples are shown. **(D)** Quantification of plots from panel B. DMSO treated samples are shown in blue and cytochalasin D treated samples are shown in orange. **(E)** Quantification of plots from panel C. **(F)** Representative images of the populations shown in panel C. Amoebae are shown in green, cell nuclei are shown in blue, and biotin is shown in magenta. Arrows indicate patches of transferred biotin. Whole human cells with stained nuclei are marked with asterisks. n=6 from 3 independent experiments.

### Protection from human serum lysis is dependent on contact with human cells

We next asked if protection from serum lysis required direct contact between amoebae and human cells, in order to determine if protection is a consequence of trogocytosis, or if protection could be acquired through secretion of proteins or release of exosomes by human cells. Amoebae and human cells were coincubated in transwell dishes, with or without direct contact (Fig 4A). After incubation, cells from the lower chambers were exposed to human serum and imaging flow cytometry was used to measure amoeba viability. Human cells were not able to pass through transwell membranes (Fig 4B). Protection from complement lysis occurred only when amoebae and human cells were incubated together in the same chamber of the transwell, but not when they were separated (Fig 4C). Protection from human serum thus required direct contact between amoebae and human cells, supporting a requirement for trogocytosis in the acquisition of protection.

**Fig 3.**
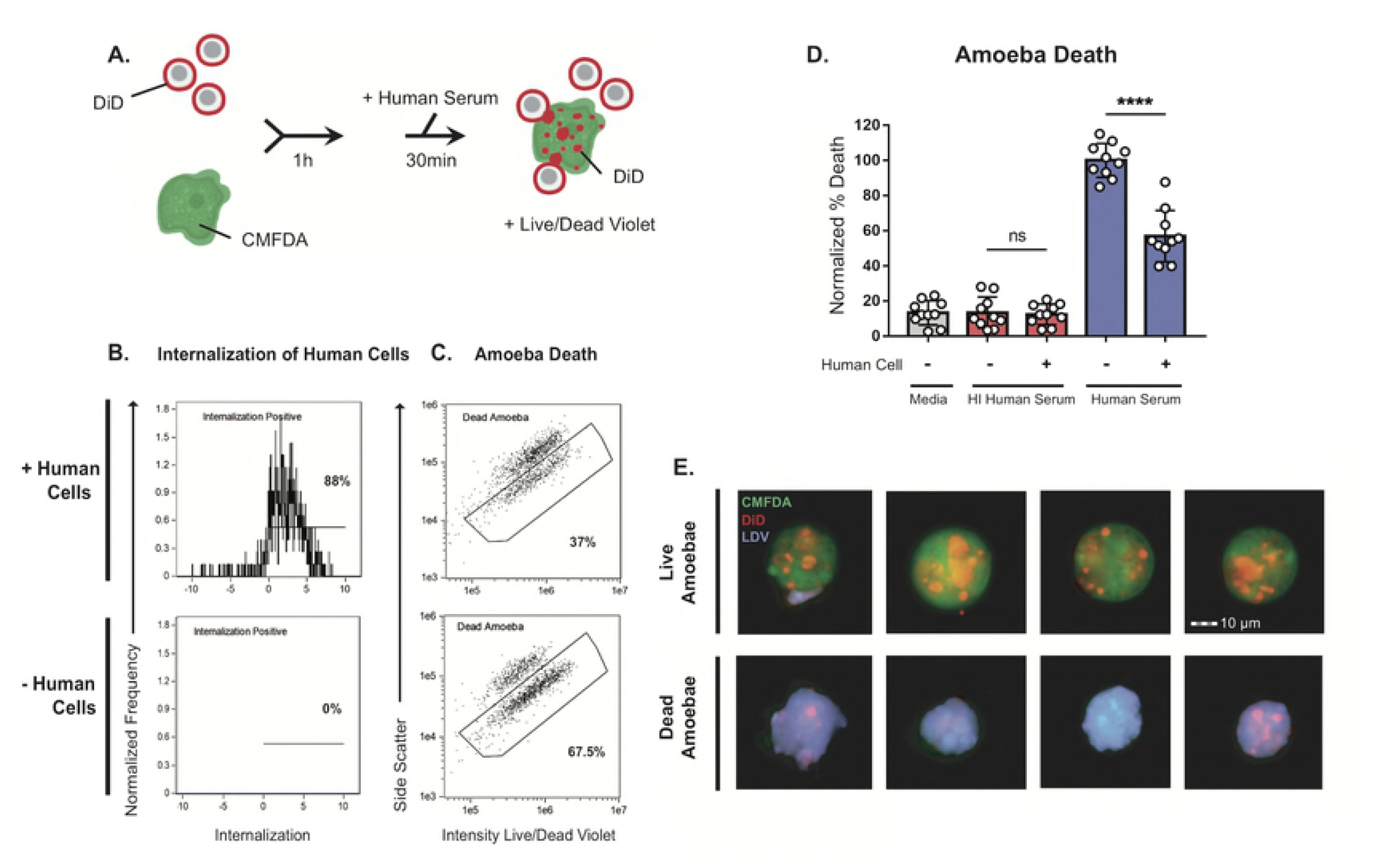
Interaction with human cells leads to protection from lysis by human serum. **(A)** CMFDA-labeled amoebae were incubated alone or in the presence of DiD-labeled human cells for 1 hour. Cells were then exposed to either active human serum, heat-inactivated human serum, or M199s medium. Following exposure to serum, samples were stained with Live/Dead Violet and viability was quantified using imaging flow cytometry, with 10,000 images collected for each sample. **(B)** Representative plots showing internalization of human cells in each condition. **(C)** Representative plots comparing amoebic death in the active serum, and heat-inactivated serum conditions. **(D)** Quantification of amoebic death for all experimental conditions. Cells exposed to M199s medium are shown in grey, heat-inactivated human serum in red, and active human serum in blue. % Death was normalized to the amoeba alone samples that were treated with active human serum. **(E)** Representative images of live and dead amoebae from amoebae co-incubated with human cells and exposed to active human serum. Amoebae are shown in green, human cells in red, and dead cells in violet. n=10 from 5 independent experiments.

**Fig 4.**
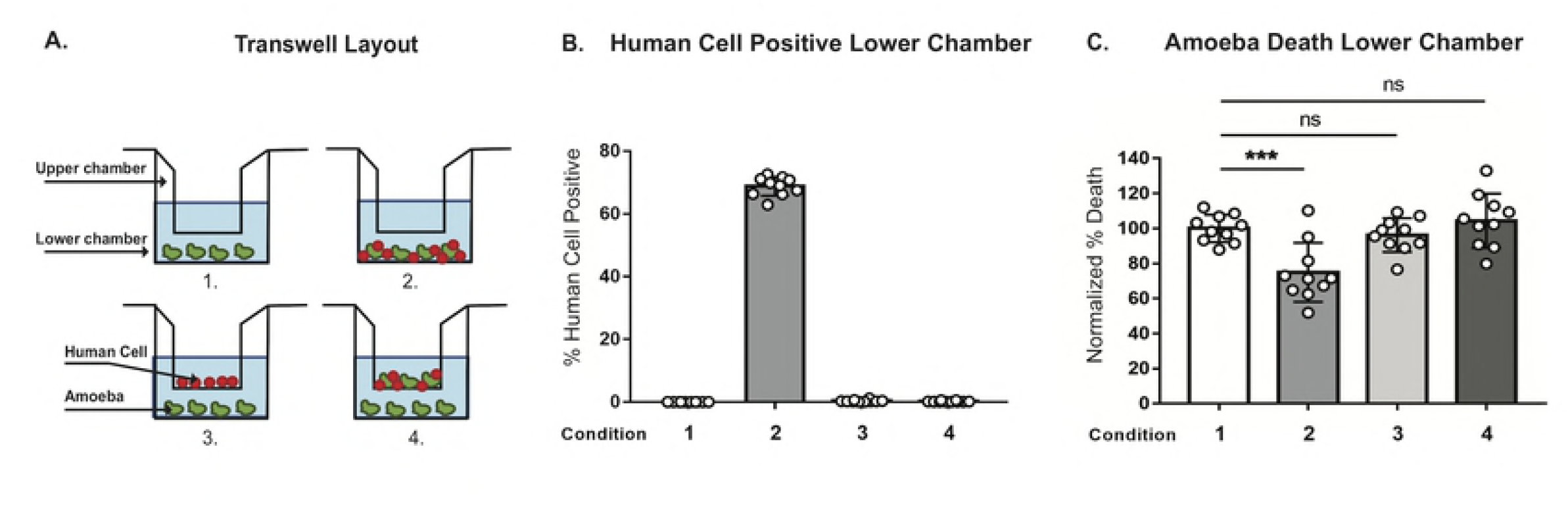
Protection from human serum lysis is dependent on contact with human cells. **(A)** Depiction of each transwell condition used in panels B-C. CMFDA-labeled amoebae and DiD-labeled human cells were incubated alone, together or separately in four different transwell conditions. Condition 1: amoebae alone in the lower chamber; condition 2: amoebae and human cells together in the lower chamber; condition 3: human cells in the upper chamber and amoebae in the lower chamber; and condition 4: amoebae and human cells together in the upper chamber and amoebae in the lower chamber. Cells were co-incubated in transwells for 1 hour and then cells from the lower chambers were harvested, exposed to human serum and analyzed. Viability was assessed using Live/Dead Violet dye and imaging flow cytometry **(B)** Quantification of human cell positive amoebae in conditions 1-4. **(C)** Quantification of amoebic death in conditions 1-4 from panel A. % Death was normalized to amoebae alone (condition 1). n=10 from 5 independent experiments.

### Protection from human serum requires actin

Since amoebic trogocytosis requires actin rearrangement [19], and acquisition and display of human cell membrane proteins requires actin (Fig 2), we next asked if treatment with cytochalasin D would also abrogate protection from human serum. Amoebae were treated with cytochalasin D, incubated in the presence or absence of human cells, and then exposed to human serum. Imaging flow cytometry was used to simultaneously measure trogocytosis (Fig 5A) and amoeba viability (Fig 5B). Amoebae that were treated with cytochalasin D were impaired in their ability to undergo trogocytosis and were not protected from serum lysis after co-incubation with human cells. Actin rearrangements are thus required for subsequent protection from lysis by human serum.

**Fig 5.**
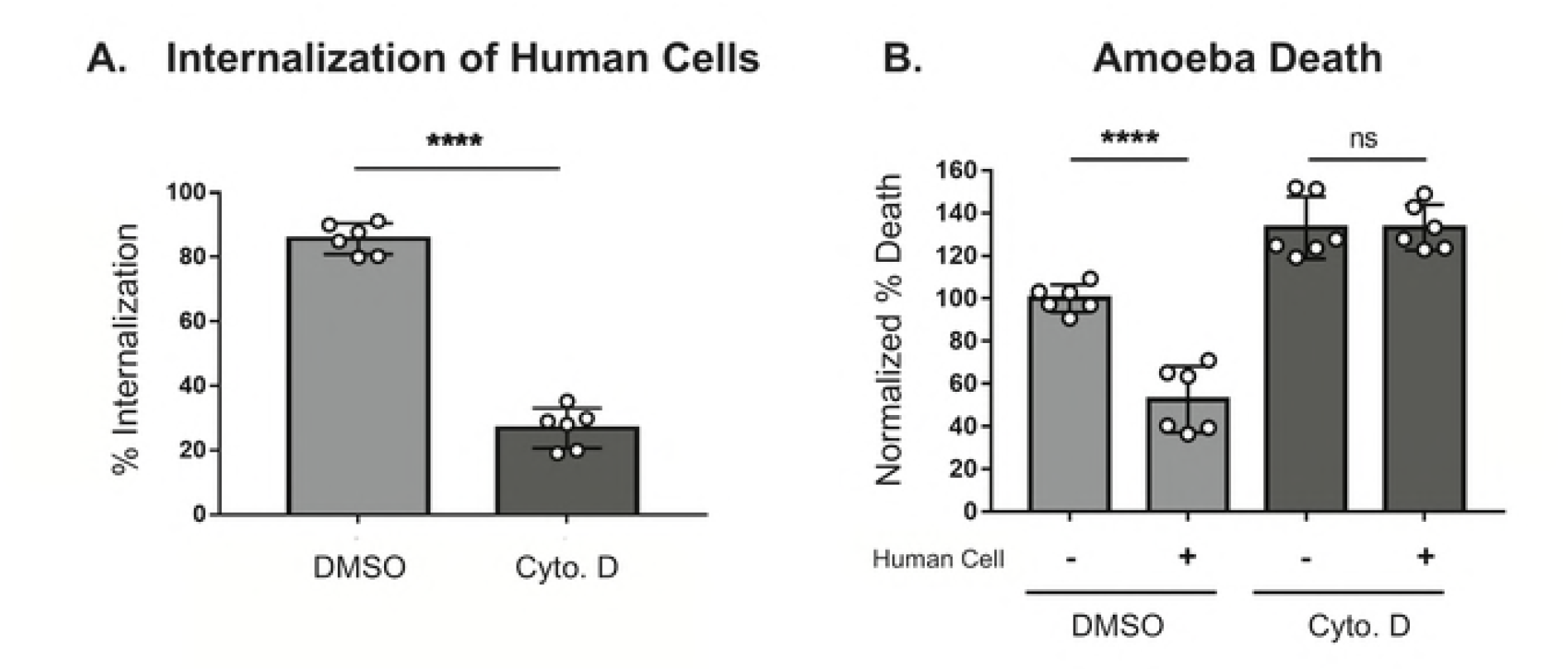
Protection from human serum is actin-dependent. CMFDA labeled amoebae were incubated alone or in the presence of DiD-labeled human cells for 1 hour and then exposed to active human serum. Samples were then stained with Live/Dead Violet viability dye and analyzed by imaging flow cytometry. **(A)** Amoebae were either pretreated with cytochalasin D (light grey) or DMSO (dark grey) for 1 hour. Quantification of internalization of human cells. **(B)** Quantification of amoebic death is shown. % Death has been normalized to the amoeba alone DMSO-treated samples. n=6 from 3 independent experiments.

### Protection occurs after trogocytosis and does not occur after phagocytosis

To ask if protection from human serum specifically occurs after trogocytosis, or if any form of ingestion leads to protection from serum, we compared amoebae that had undergone trogocytosis versus those that had undergone phagocytosis. We previously showed that amoebae undergo trogocytosis of live human cells, and in contrast, undergo phagocytosis of pre-killed human cells [19]. Therefore, we asked if phagocytosis of pre-killed cells could also provide protection from complement lysis. Human cells were pretreated with staurosporine to induce apoptosis (Fig 6A). Amoebae were co-incubated with live or pre-killed human cells or incubated in the absence of human cells. Amoebae that had undergone trogocytosis or phagocytosis ingested a similar amount of human cell material (Fig 6B), however, amoebae were only protected from lysis by human serum after undergoing trogocytosis (Fig 6C). Therefore, protection from lysis by human serum occurs specifically after trogocytosis of live cells and not phagocytosis of pre-killed cells.

**Fig 6.**
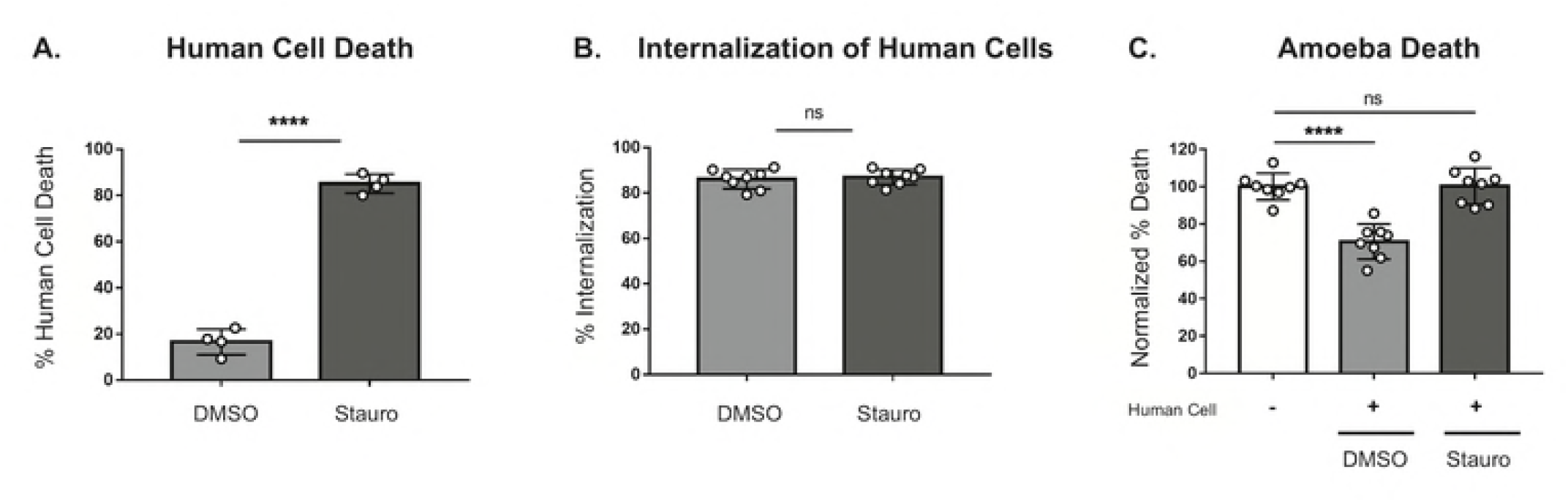
Protection requires trogocytosis but not phagocytosis of human cells. **(A)** Human cells were pretreated with staurosporine (dark grey) or DMSO (light grey). The human cell viability before co-incubation is shown. **(B)** Quantification of human cell internalization by amoebae. **(C)** Quantification of amoebic death. % Death was normalized to the amoebae alone samples. n=8 from 4 independent experiments.

To further distinguish between requirements for trogocytosis and phagocytosis, we next tested knockdown mutants deficient in rhomboid protease 1 (*Eh*ROM1) (EHI_197460), a rhomboid protease with roles in attachment and ingestion [39,40]. *Eh*ROM1 mutants have been shown to be deficient in both phagocytosis and pinocytosis, as well as attachment to live cells [39,40]. Furthermore, it has been shown that silencing of *Eh*ROM1 does not change susceptibility to serum lysis, making these mutants an ideal tool for testing the effects conferred by ingestion of human cells [39,40]. We generated stable transfectants knocked down for expression of *Eh*ROM1 (Fig 7A). *Eh*ROM1 knockdown mutants were deficient in attachment to healthy human cells (Fig 7B-C), consistent with previous studies [40]. Also consistent with previous studies, *Eh*ROM1 mutant amoebae incubated alone were not more susceptible to serum lysis then control amoebae (S5B Fig). Using imaging flow cytometry, we assayed *Eh*ROM1 mutants for a trogocytosis defect for the first time and found that they did not exhibit a defect in trogocytosis of live human cells (Fig 7D, S6 Fig). Consistent with previous studies [40], we found that *Eh*ROM1 mutants were defective in phagocytosis of pre-killed human cells (Fig 7E). After co-incubation with live human cells, *Eh*ROM1 mutants were no more or less protected from lysis by human serum than control amoebae (Fig 7F, S5 Fig). Therefore, a mutant deficient in phagocytosis does not exhibit a difference in protection from serum, further supporting that phagocytosis is not involved in resistance to lysis by human serum. Moreover, resistance to lysis by human serum is not associated with simple attachment to human cells, since *Eh*ROM1 mutants are impaired in binding to live human cells but still exhibit no difference in resistance to human serum. Together, these finding further underscore that protection from lysis by human serum is associated with trogocytosis, not phagocytosis.

**Fig 7.**
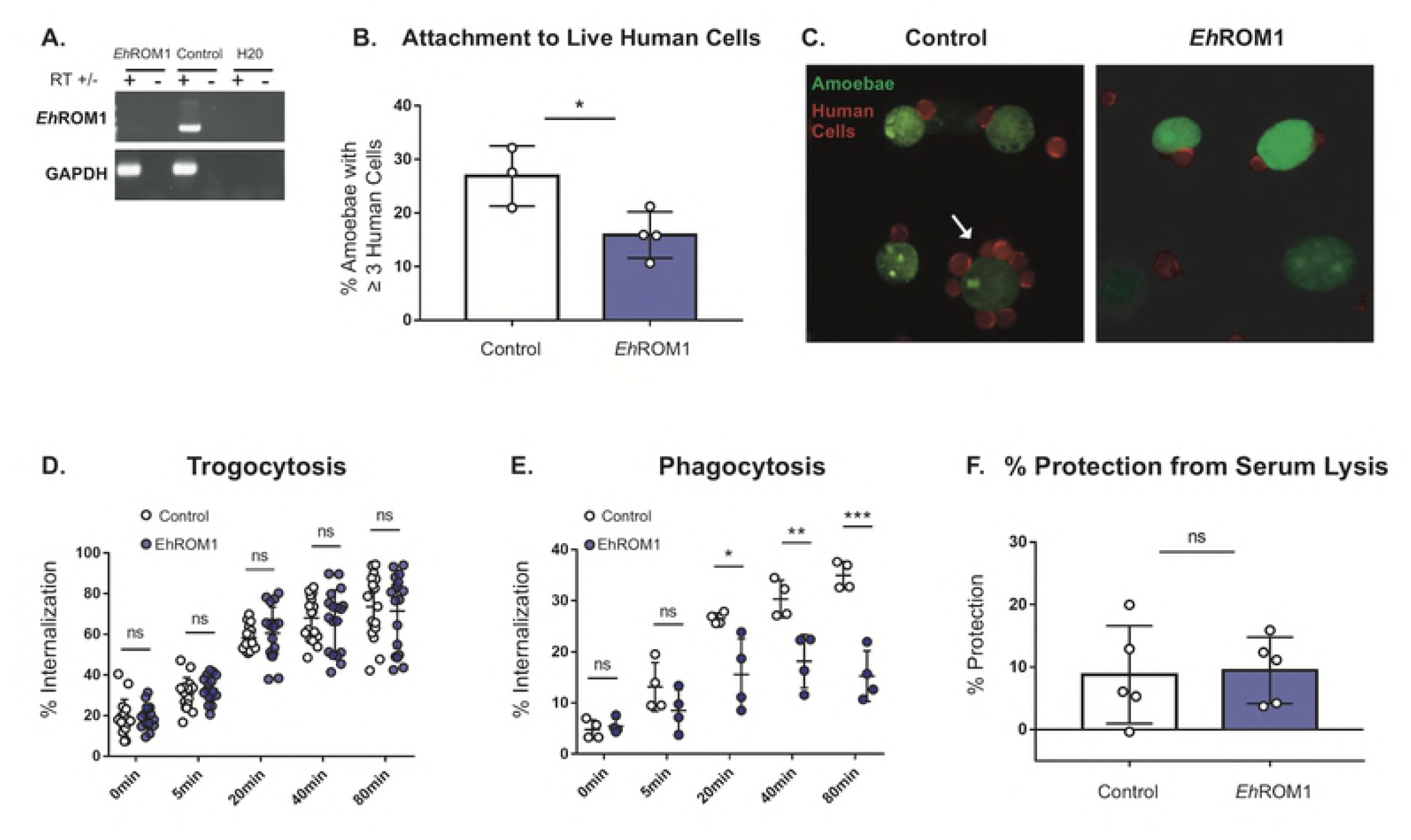
*Eh*ROM1 knockdown mutants defective in phagocytosis but not trogocytosis are protected from serum lysis. Amoebae were stably transfected with an *Eh*ROM1 knockdown plasmid (EhROM1) or vector control plasmid (Control). **(A)** Silencing of *Eh*ROM1 was verified by using RT-PCR. Reverse transcriptase (RT) was included (+), or omitted (-) as a control. GAPDH was used to control for loading. **(B)** *Eh*ROM1 and vector control transfectants were incubated on ice with live human cells for 1 hour, then fixed and analyzed using confocal microscopy. The percentage of amoebae with 3 or more attached human cells for each condition is displayed; vector control is shown with open bars and the *Eh*ROM1 knockdown mutant is shown with blue bars. n=4 replicates from 2 independent experiments. 20 images were collected per slide and 195-252 individual amoebae were counted per condition. **(C)** Representative images from panel B. Amoebae are shown in green and human cells are shown in red. Arrow indicates an amoeba with a rosette of attached human cells. **(D)** CMFDA-labeled *Eh*ROM1 knockdown mutants (blue circles) or vector control (open circles) transfectants were incubated alone or in the presence of live DiD-labeled human cells for 0, 5, 20, 40 or 80 minutes. Internalization of human cell material was quantified using imaging flow cytometry. n=20 from 10 independent experiments. **(E)** CMFDA-labeled *Eh*ROM1 knockdown mutants (blue circles) or vector control (open circles) transfectants were incubated alone or in the presence of heat-killed CTDR-labeled human cells for 0, 5, 20, 40 or 80 minutes. Internalization of human cell material was quantified using imaging flow cytometry. n=4 from 2 independent experiments. **(F)** *Eh*ROM1 (blue bar) or vector control (open bar) amoebae were co-incubated with live human cells for 1 hour, and then exposed to human serum. Viability was assessed using Live/Dead Violet dye and imaging flow cytometry. % protection was calculated by subtracting the total lysis of amoebae co-incubated with human cells from the total lysis of amoebae incubated alone. n= 9-10 from 5 independent experiments. Protection data displays means of 2 replicates from all 5 experiments.

Collectively, these results support a new model of immune evasion in which amoebae perform trogocytosis on live human cells and through trogocytosis, acquire and display human cell membrane proteins. Display of human cell membrane proteins then leads to protection from human serum, most likely by inhibiting complement-mediated lysis (Fig 8).

**Fig 8.**
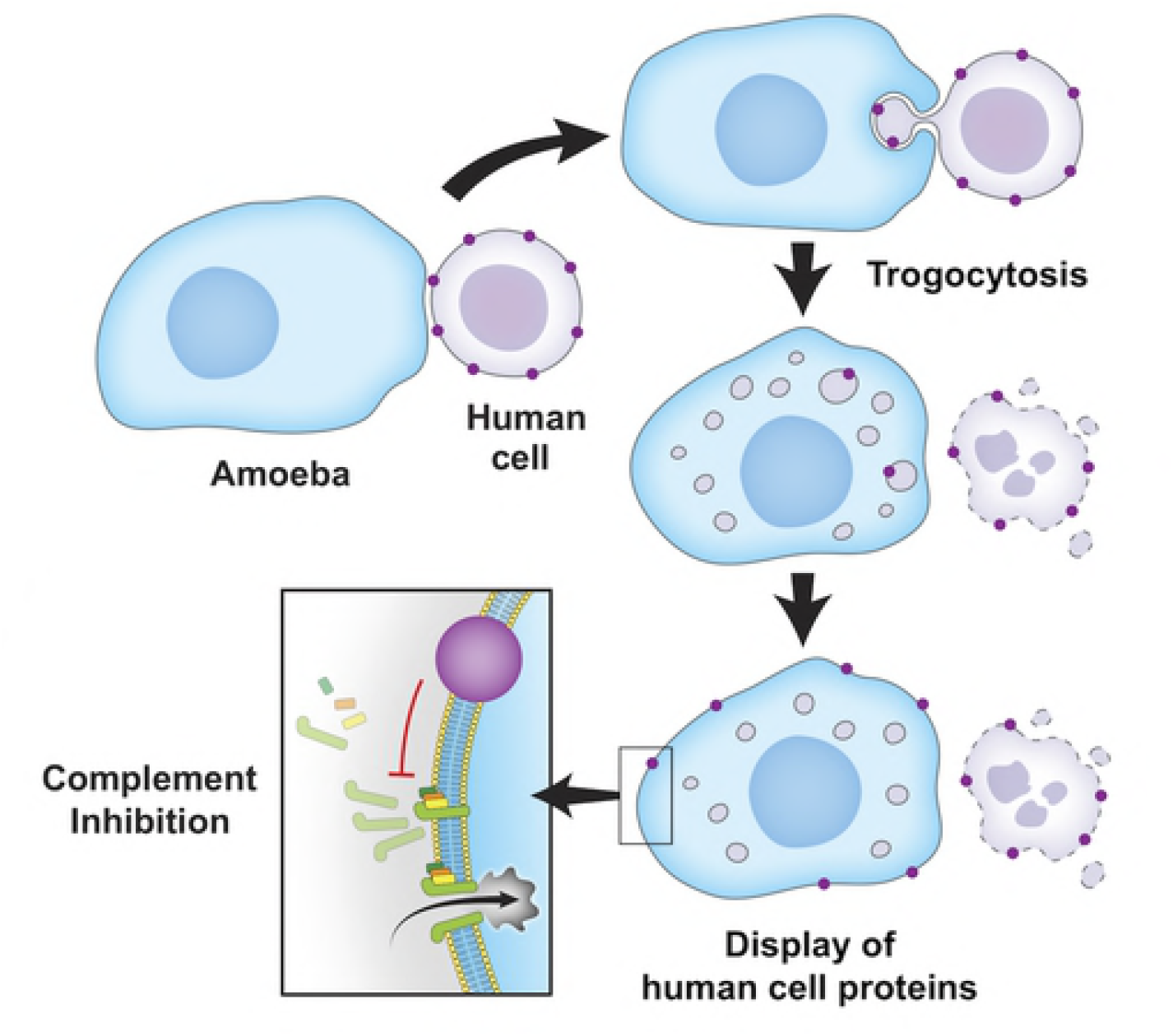
Proposed model of protection from serum lysis. Amoebae encounter live human cells while invading the intestine or disseminating in the blood stream and perform trogocytosis. Trogocytosis leads to acquisition and display of human cell membrane proteins on the amoebae surface. Display of human cell proteins then protects the amoebae from lysis in the blood by inhibiting the complement cascade.

## Discussion

In this study, confocal microscopy and imaging flow cytometry experiments revealed that amoebae acquire and display human cell membrane proteins. This process is actin-dependent and is associated with resistance to lysis by human serum. Protection from lysis by human serum requires direct contact between amoebae and human cells, is actin-dependent, and is specifically associated with trogocytosis, not phagocytosis. Collectively, these data suggest that amoebae acquire and display human cell membrane proteins through trogocytosis, and that this leads to protection from lysis by human serum complement.

Complement resistance by amoebae appears to be relevant to invasive disease. Once amoebae have invaded intestinal tissue, they can spread from the intestine to the liver through the portal vein [41], and they can ingest erythrocytes [42], thus they are capable of surviving in the bloodstream. A study that depleted complement by using cobra venom factor in the hamster model of amoebic liver abscess, found that loss of complement was correlated with greater severity of liver lesions [43]. Additionally, it was found that serum from women was more effective in killing amoebae then serum from men, and men are known to be more susceptible to invasive amoebiasis [44]. Furthermore, pathogenic amoebae have been shown to resist complement. *E. histolytica* appears to evade complement deposition, while the closely related nonpathogenic species *Entamoeba dispar* does not [45]. Similarly, amoebae isolated from patients with invasive infection resist complement, while strains isolated from asymptomatic patients are complement-sensitive [46].

Previous studies have hinted that amoebae become more resistant to complement after interacting with host cells or tissues, and that complement resistance involves proteins on the amoeba surface. Consistent with our findings, it has previously been demonstrated that amoebae that were made resistant to complement lysis by hamster liver passage, lost resistance after treatment with trypsin [38], suggesting that complement-resistance is associated with proteins on the amoeba surface. It has also been shown that amoebae acquire serum resistance after ingestion of live human erythrocytes, and that resistant amoebae stained positive with antiserum directed to erythrocyte membrane antigens [36]. Though this previous study described ingestion of erythrocytes as erythrophagocytosis, we now know that amoebae are also capable of performing trogocytosis on live erythrocytes [19]. In older literature, amoebae were also seen to ingest bites of erythrocytes in a process that was termed microphagocytosis [47]. Therefore, we propose a model in which invasive amoebae are able to evade complement detection in the blood by trogocytosis of human cells and display of human cell membrane proteins.

Other mechanisms of complement resistance in *E. histolytica* have been described such as mimicry of the complement regulatory protein CD59 [48,49], an inhibitor of the membrane attack complex (MAC). Amoebic cysteine proteinases play a role in cleavage of complement components [50–52]. It has also been reported that amoebae are made temporarily resistant to complement lysis through treatment with increasing doses of heat-inactivated human serum, though the mechanism remains unclear [37,53], and it was recently found that amoebae do not develop resistance to serum from rats by this method [54]. As the percentage of amoebae lysed after exposure to human serum in our assays never reached 100%, even in conditions where amoebae were incubated alone, it is likely that multiple factors contribute to complement resistance in *E. histolytica.* We propose that acquisition of human cell membrane proteins is one of the mechanisms by which amoebae evade lysis by complement.

It will be of great interest to determine which proteins are transferred and displayed on the amoeba surface. It is possible that complement regulatory proteins such as CD55 or CD46 are displayed by amoebae and that this directly promotes resistance to complement lysis. Displayed human cell membrane proteins may also bind to soluble factors in human serum, such as factor H. It is notable that acquired human cell membrane proteins do not have an even distribution on the amoeba surface, and instead appear in patches. This was similar for both biotin-streptavidin and MHC-I staining. In mammalian immune cells, similar patchy localization of acquired membrane proteins has been seen, and this was seen with biotin-streptavidin staining, fluorescently-tagged proteins, and immunofluorescence [24,28]. It is not clear if the acquired membrane proteins are present in lipid microdomains (*e.g.,* lipid rafts), or are in clusters. It is also possible that while patchy foci of acquired membrane proteins are clearly seen, these proteins may also be found throughout the membrane at lower concentrations below the limit of detection. In any case, the distribution of human cell proteins appears sufficient to confer protection from complement.

With the discoveries of amoebic trogocytosis and display of human cell membrane proteins, a new paradigm for amoeba-human cell interactions is emerging. We previously showed that when amoebae kill cells, they do not ingest dead cell corpses [19]. Prior to this, amoebae were thought to fully ingest the corpses of the cells they had killed [5,55,56]. Now, with the discovery of acquisition and display of human cell membrane proteins, together with the lack of ingestion of cell corpses, a different paradigm is emerging. It is possible that rather than acquiring nutrition by killing and ingesting entire cells, amoebae nibble and acquire membrane proteins that contribute to immune evasion. Invasive disease involves survival of amoebae in blood vessels. Since trogocytosis contributes to tissue invasion [19], it is possible that amoebae acquire human cell membrane proteins as they invade the intestine. Amoebae would then be equipped to survive in the bloodstream and to spread to other tissues. Moreover, since there is the potential for a variety of human cell proteins to be displayed, display of human cell proteins may impact host-amoeba interactions in many ways.

Display of human cell proteins acquired during trogocytosis is a novel strategy for immune evasion by a pathogen. Since other microbial eukaryotes use trogocytosis for cell killing, including *N. fowleri,* there is the potential for display of acquired membrane proteins to apply to the pathogenesis of other infections. Furthermore, our studies extend acquisition and display of membrane proteins beyond mammalian immune cells, suggesting that this may be a fundamental feature of eukaryotic trogocytosis. How membrane proteins are acquired and displayed by immune cells during trogocytosis is not well understood in immune cells and, to our knowledge, the underlying mechanism is not under investigation. Thus, ongoing studies in amoebae may shed light on acquisition and display of membrane proteins during trogocytosis.

In summary, we have shown that amoebae display human cell membrane proteins on their surface and are protected from lysis by human serum after trogocytosis of live human cells. We propose a new model of immune evasion by *E. histolytica,* whereby amoebae survive complement attack in the blood through trogocytosis of human cells and display of human cell membrane proteins. This work broadens our understanding of trogocytosis as a conserved feature of eukaryotic biology, as well as our understanding of the pathogenesis of amoebiasis.

## Materials and Methods

### Cell culture

HM1: IMSS (ATCC) *E. histolytica* trophozoites (amoebae) were cultured at 35°C in Trypticase-Yeast Extract-Iron-Serum (TYI-S-33) media supplemented with 80 Units/mL penicillin and 80 μg/mL streptomycin (Gibco), 2.3% Diamond Vitamin Tween 80 Solution 40x (Sigma-Aldrich) and 15% heat-inactivated adult bovine serum (Gemini Bio-Products). Amoebae were harvested when tissue culture flasks reached 80-100% confluency and then resuspended in M199s media (Gibco medium M199 with Earle’s Salts, L-Glutamine, 2.2 g/L Sodium Bicarbonate and without Phenol Red) supplemented with 5.7 mM L-cysteine (Sigma-Aldrich), 25 mM HEPES (Sigma-Aldrich) and 0.5% bovine serum albumin (Gemini Bio-Products).

Human Jurkat T cells from ATCC (Clone E6-1) were cultured at 37°C and 5% CO_2_ in RPMI Medium 1640 (Gibco RPMI with L-Glutamine and without Phenol Red) supplemented with 10 mM HEPES (Affymetrix), 100 Units/mL penicillin and 100 μg/mL streptomycin (Gibco) and 10% heat-inactivated fetal bovine serum (Gibco). human cells were harvested between 5×10^5^ and 2×10^6^ cells/ml and resuspended in M199s media.

### Generation of *Eh*ROM1 mutants

The *Eh*ROM1 silencing construct, made from a pEhEx plasmid backbone, was generated by Morf *et al.* as described in [57]. The construct contained 132 base pairs of the trigger gene EHI_048600 fused to the first 537 base pairs of *Eh*ROM1 (EHI_197460). Amoebae were transfected with 20 μg of the *Eh*ROM1 silencing construct using Attractene Transfection Reagent (QIAGEN). Transfectants were then maintained under selection with Geneticin at 6 μg/ml. Clonal lines were generated by limiting dilution in a 96-well plate contained in a BD GasPak EZ Pouch System (BD Biosciences), and silencing was confirmed with RT-PCR. An individual clonal line was used for all experiments. A vector control line was generated by transfection with the pEhEx-trigger construct backbone, using the same approach.

### Confocal immunofluorescence assays

Amoebae were washed and labeled in M199s with CellTracker Green CMFDA Dye (Invitrogen) at 310 ng/ml for 10 minutes at 35°C. In the biotin transfer experiments, human cells were resuspended in 1X Dulbecco’s Phosphate Buffered Saline (PBS: Sigma-Aldrich) and then biotinylated with EZ-Link Sulfo-NHS-SS-Biotin (Thermo Fisher Scientific) at 480 μg/ml in 1X PBS for 25 minutes at 4°C. 1M Tris-HCL pH 8 was added to the samples for a final concentration of 100 mM to quench the reaction. Cells were next washed in 1X PBS containing Tris-HCL pH 8 at 100 mM, and then resuspended in M199s. Amoebae and human cells were combined at a 1:5 ratio in M199s and co-incubated for 5 minutes at 35°C. Following co-incubation, cells were fixed with 4% paraformaldehyde (Electron Microscopy Sciences) for 30 minutes at room temperature and stained with an Alexa Fluor 633 streptavidin conjugate (Invitrogen) at 20 μg/ml for 1 hour at 4°C. After fixation, samples were stained with DAPI (Sigma-Aldrich) for 10 minutes at room temperature. Samples were then incubated on coverslips pre-coated with collagen (Collagen I, Rat Tail: Gibco), according to the manufacturer’s instructions, for 1 hour at room temperature and mounted on glass slides using VECTASHIELD Antifade Mounting Medium (Vector Laboratories). In some experiments, samples were incubated on Superfrost Plus Micro Slides (VWR) for 1 hour, and coverslips were then mounted with VECTASHIELD Antifade Mounting Medium. Samples were imaged on an Olympus FV1000 laser point-scanning confocal microscope or on an Intelligent Imaging Innovations Hybrid Spinning Disk Confocal-TIRF-Widefield Microscope. Images were collected from 4 independent experiments.

For the MHC class I immunofluorescence experiments, human cells were washed and resuspended in M199s but left unlabeled before co-incubation with amoebae. Amoebae and human cells were combined at a 1:5 ratio in M199s and co-incubated for 5 minutes at 35°C. Following co-incubation and fixation, samples were blocked for 1 hour in PBS-T (0.1% Tween 20 in 1X PBS) supplemented with 20% Goat Serum (Jackson Immunoresearch Labs Inc.) and 5% bovine serum albumin (Gemini Bio-Products). Samples were then washed in PBS-T and incubated overnight with an MHC class I monoclonal primary antibody (Thermo Fisher Scientific HLA-ABC Monoclonal Antibody W6/32) at 10 μg/ul, followed by washing with PBS-T and incubation with an anti-mouse Cy3 secondary antibody (Jackson Immunoresearch Labs Inc.) at 3.5 ng/ml at room temperature for 1 hour. Samples were stained with DAPI and mounted on glass slides as above. Images were collected from 4 independent experiments.

### Imaging flow cytometry immunofluorescence assays

Amoebae were resuspended in M199s media and pretreated with cytochalasin D from *Zygosporium mansonii* (Sigma-Aldrich) at 20 μM or with the equivalent volume of dimethylsulfoxide (DMSO) for 1 hour at 35°C. Cytochalasin D and DMSO were kept in the media for the duration of the experiment. Following pretreatment, amoebae were labeled with CellTracker Green CMFDA Dye (Invitrogen) at 93 ng/ml for 10 minutes at 35°C. Human cells were labeled in culture with Hoechst 33342 (Invitrogen) at 5 μg/ml for 1 hour at 37°C and then resuspended in 1X PBS. Human cells were then biotinylated with EZ-Link Sulfo-NHS-SS-Biotin (Thermo Fisher Scientific) at 480 μg/ml in 1X PBS for 25 minutes at 4°C. 100 mM Tris-HCL pH 8 was used to quench the reaction, cells were washed in 100 mM Tris-HCL pH 8 and were resuspended in M199s. Amoebae and human cells were combined at a 1:5 ratio in M199s and co-incubated for 5 minutes at 35°C. After coincubation, samples were immediately placed on ice to halt ingestion, stained with an Alexa Fluor 633 streptavidin conjugate (Invitrogen) at 20 μg/ml for 1 hour at 4°C and fixed with 4% paraformaldehyde (Electron Microscopy Sciences) for 30 minutes at room temperature. Fixed samples were resuspended in 1X PBS and run on an Amnis ImageStreamX Mark II. 10,000 events per sample were collected from 6 repeats across three independent experiments.

### Serum lysis assays

Amoebae were washed and labeled in M199s with CellTracker Green CMFDA Dye (Invitrogen) at 93 ng/ml for 10 minutes at 35°C. Human cells were washed and labeled in M199s with Diic18(5)-Ds [1,1-Dioctadecyl-3,3,3,3-Tetramethylindodicarbocyanine-5,5-Disulfonic Acid] (DiD: Assay Biotech) at 21 μg/ml for 5 minutes at 37°C and 10 minutes at 4°C. After washing with M199s, amoebae and human cells were combined at a 1:5 ratio in M199s and co-incubated for 1 hour at 35°C, or amoebae were incubated in the same conditions in the absence of human cells. Next, cells were pelleted at 400 × *g* for 8 minutes and were resuspended in 100% normal human serum (Pooled Normal Human Complement Serum, Innovative Research Inc.), heat-inactivated human serum (inactivated at 56°C for 30 minutes), or M199s. Serum/media was supplemented with 150 μM CaCl_2_ and 150 μM MgCl_2_ (Fig. S2). Next, cells were incubated for 30 minutes at 35°C. Cells were then washed and resuspended in M199s media and incubated with LIVE/DEAD Fixable Violet Dead Cell Stain (Invitrogen) that was prepared according to the manufacturer’s instructions, at 4 μl/ml for 30 minutes on ice. Next, samples were fixed with 4% paraformaldehyde (Electron Microscopy Sciences) for 30 minutes at room temperature. Fixed samples were pelleted and resuspended in 1X PBS, then run on an Amnis ImageStreamX Mark II. 10,000 events per sample were collected.

In the cytochalasin D experiments, amoebae were pretreated with cytochalasin D from *Zygosporium mansonii* (Sigma-Aldrich) at 20 μM or an equivalent volume of DMSO for 1 hour at 35°C. Cytochalasin D/DMSO was kept in the media for the duration of the experiment. In experiments where amoebae ingested live or pre-killed cells, human cells were pretreated in culture with staurosporine from *Streptomyces* sp. (Sigma-Aldrich) at 1 μM or with the equivalent volume of DMSO overnight at 37°C. Human cells were then washed and suspended in M199s media and labeled with CellTracker Deep Red (CTDR) (Invitrogen) at 1 μM for 30 minutes at 37°C. In transwell assays, amoebae and human cells were incubated together at a 1:5 ratio or separately in 12 mm transwells with 3.0 μm pore, 10 μm thick polycarbonate membrane inserts (Corning). In experiments using EhROM1 knockdown, stably transfected *Eh*ROM1 clonal mutants were compared to mutants that contained a pEhEx-trigger backbone vector control construct.

### Ingestion assays

In trogocytosis assays, CMFDA labeled transfectants were incubated alone or in the presence of live DiD-labeled Jurkat cells for 0, 5, 20, 40 or 80 minutes. Samples were then labeled with Live/Dead Violet and fixed with 4% paraformaldehyde. Internalization of human cell material was quantified using imaging flow cytometry. In phagocytosis assays, human cells were heat-killed at 60°C for 40 minutes and were labeled with CTDR and Hoechst prior to incubation with CMFDA labeled amoebae.

### Attachment assay

CMFDA labeled amoebae were combined with CTDR labeled live human cells at a 1:5 ratio, centrifuged at 150 × g for 5 minutes 4°C, and incubated on ice for 1 hour. Samples were then fixed with 4% paraformaldehyde. Samples were incubated on Superfrost Plus Micro Slides (VWR) for 1 hour, coverslips were mounted with VECTASHIELD Antifade Mounting Medium and slides were imaged on an Intelligent Imaging Innovations Hybrid Spinning Disk Confocal-TIRF-Widefield Microscope. 20 images were collected per slide. Amoebae with 3 or more attached human cells were scored as attachment positive. Image collection and scoring were performed in a blinded manner.

### Imaging flow cytometry analysis

Samples were run on an Amnis ImageStreamX Mark II and 10,000 events were collected per sample. Data were analyzed using Amnis IDEAS software. Samples were gated on focused cells, single amoebae, amoebae that had come in contact with human cells, and amoebae that had internalized human material. From the single amoebae gate, amoebic death was quantified by plotting intensity of LIVE/DEAD Violet against side scatter and gating on LIVE/DEAD Violet positive cells (see Fig. S3).

In the biotin transfer experiment, single amoebae were divided into Hoechst high and Hoechst low populations in order to isolate single amoebae with and without human cells. Overlap of biotin with CMFDA labeled amoebae was plotted and biotin positive cells were selected from both Hoechst high and low populations (see Fig. S1).

In the trogocytosis and phagocytosis assays, focused cells were gated from total collected events. Next, single cells were gated, and then single amoebae were gated. Amoebae positive for human cells were gated and internalization of human cells was measured. (see Fig. S5)

### Statistical analysis

All statistical analysis was performed using GraphPad Prism. All data plots display means and standard deviation values. Data were statistically analyzed using a student’s unpaired t-test (ns = P>0.05, * = P ≤ 0.05, ** = P ≤ 0.01, *** = P ≤ 0.001, **** = P ≤ 0.0001).

## Acknowledgements

We thank the MCB Light Microscopy Imaging Facility at UC Davis for technical assistance. We thank Dr. Stephen McSorley and the members of our laboratory for helpful discussions.

## Supporting Information

**S1 Fig: Gating strategy used to quantify transferred biotin.**

Gating strategy used to quantify biotin-positive amoebae. Focused cells were gated from total collected events. Next, single cells were gated, and then single amoebae were gated. Single amoebae were divided into Hoechst high and Hoechst low populations to identify images with and without human cell nuclei. Finally, biotin-positive amoebae were gated on from images with and without human cell nuclei.

**S2 Fig: Optimization of complement assay.**

The ability of un-supplemented human serum from different vendors to lyse amoebae was tested at various concentrations for 30 minutes, 1 hour, and 2 hours at 35°C. Samples were labeled with the viability dye Live/Dead Violet and % amoeba death was quantified using imaging flow cytometry. % amoeba death was not normalized. **(A)** Sigma Male AB Serum. Note, serum was stored at −20°C instead of −80°C. **(B)** Sigma Complement Sera Human Lyophilized Powder. **(C)** Innovative Research Pooled Normal Human Complement Serum. **(D)** Valley Biomedical Human Complement (Serum). **(E-F)** Lysis of increasing concentration of serum from Innovative Research and Valley Biomedical was tested with the addition of 150 μM CaCl_2_ and 150 μM MgCl_2_ for 1 hour at 35°C.

**S3 Fig: Serum lysis assay gating strategy.**

Gating strategy used in the serum lysis assay. Focused cells were gated from total collected events. Next, focused events were dived into gates that either contained debris and human cells, or single amoebae. Single amoebae positive for human cells were gated and then internalization of human cells was measured. % of dead amoebae was gated from single amoebae.

**S4 Fig: Non-normalized data shown from Fig 3.**

Non-normalized data from the serum lysis assay shown in Figure 3. Amoebic lysis was variable and fell in to two groups, Low lysis **(A)** and **(B)** high lysis. This variability in lysis was associated with how the human serum was stored and thawed. The highest lysis was achieved with serum stored at −80°C and rapidly thawed at 37 °C, leaving intact ice pellets, and then thawed to completion at room temperature. Lower lysis was achieved with serum stored at −20°C and thawed to completion at 37 °C. **(C)** Lysis from all data non-normalized. **(D)** Lysis from all data normalized to the amoeba incubated alone condition that was exposed to active human serum.

**S5 Fig: Internalization of human cells and amoebic death from the serum lysis assay in Fig 7.**

Additional data from the serum lysis assay used in Figure 7. **(A)** Internalization of human cells by vector control transfectants (open bar) or *Eh*ROM1 (blue bar) knockdown mutants. **(B)** % of normalized amoeba death in the conditions where amoebae were incubated alone. **(C)** Non-normalized amoebic death from all conditions.

**S6 Fig: Gating strategy used in trogocytosis and phagocytosis assays.**

**(A)** Gating strategy used in the trogocytosis and phagocytosis assays shown in Figure 7. Shown are example data from a trogocytosis assay, where CMFDA-labeled amoebae were incubated with live DiD-labeled human cells. For phagocytosis assays, CMFDA-labeled amoebae were incubated with CTDR-labeled heat-killed human cells. Focused cells were gated from total collected events. Next, single cells were gated, and then single amoebae were gated. Amoebae positive for human cells were gated and internalization of human cells was measured. **(B)** Example data from a trogocytosis assay, with representative plots from the vector control condition showing internalization of human cells over time as well as representative images collected at each time point.

## References

1. Haque R, Mondal D, Kirkpatrick BD, Akther S, Farr BM, Sack RB, et al. Epidemiologic and clinical characteristics of acute diarrhea with emphasis on Entamoeba histolytica infections in preschool children in an urban slum of Dhaka, Bangladesh. Am J Trop Med Hyg. 2003;69: 398–405.

2. Speich B, Croll D, Fürst T, Utzinger J, Keiser J. Effect of sanitation and water treatment on intestinal protozoa infection: a systematic review and meta-analysis. The Lancet Infectious Diseases. 2016;16: 87–99. doi:10.1016/S1473-3099(15)00349-7

3. Petri WA, Mondal D, Peterson KM, Duggal P, Haque R. Association of malnutrition with amebiasis. Nutr Rev. 2009;67: S207–S215. doi:10.1111/j.1753-4887.2009.00242.x

4. Gilchrist CA, Petri SE, Schneider BN, Reichman DJ, Jiang N, Begum S, et al. Role of the Gut Microbiota of Children in Diarrhea Due to the Protozoan Parasite Entamoeba histolytica. J Infect Dis. 2016;213: 1579–1585. doi:10.1093/infdis/jiv772

5. Ralston KS, Petri WA. Tissue destruction and invasion by Entamoeba histolytica. Trends Parasitol. 2011;27: 254–263. doi:10.1016/j.pt.2011.02.006

6. Ravdin JI, Croft BY, Guerrant RL. Cytopathogenic mechanisms of Entamoeba histolytica. J Exp Med. 1980;152: 377–390.

7. Ravdin JI, Guerrant RL. Studies on the cytopathogenicity of Entamoeba histolytica. Arch Invest Med (Mex). 1980;11: 123–128.

8. Ravdin JI, Guerrant RL. Role of Adherence in Cytopathogenic Mechanisms of Entamoeba Histolytica. J Clin Invest. 1981;68: 1305–1313.

9. Saffer LD, Petri WA. Role of the galactose lectin of Entamoeba histolytica in adherence-dependent killing of mammalian cells. Infect Immun. 1991;59: 4681–4683.

10. Thibeaux R, Dufour A, Roux P, Bernier M, Baglin A-C, Frileux P, et al. Newly visualized fibrillar collagen scaffolds dictate Entamoeba histolytica invasion route in the human colon. Cell Microbiol. 2012;14: 609–621. doi:10.1111/j.1462-5822.2012.01752.x

11. Hellberg A, Nickel R, Lotter H, Tannich E, Bruchhaus I. Overexpression of cysteine proteinase 2 in Entamoeba histolytica or Entamoeba dispar increases amoeba-induced monolayer destruction in vitro but does not augment amoebic liver abscess formation in gerbils. Cell Microbiol. 2001;3: 13–20.

12. Keene WE, Petitt MG, Allen S, McKerrow JH. The major neutral proteinase of Entamoeba histolytica. J Exp Med. 1986;163: 536–549.

13. Lidell ME, Moncada DM, Chadee K, Hansson GC. Entamoeba histolytica cysteine proteases cleave the MUC2 mucin in its C-terminal domain and dissolve the protective colonic mucus gel. Proc Natl Acad Sci USA. 2006;103: 9298–9303. doi:10.1073/pnas.0600623103

14. Bracha R, Nuchamowitz Y, Leippe M, Mirelman D. Antisense inhibition of amoebapore expression in Entamoeba histolytica causes a decrease in amoebic virulence. Mol Microbiol. 1999;34: 463–472.

15. Bracha R, Nuchamowitz Y, Mirelman D. Transcriptional silencing of an amoebapore gene in Entamoeba histolytica: molecular analysis and effect on pathogenicity. Eukaryotic Cell. 2003;2: 295–305.

16. Leippe M, Andrä J, Müller-Eberhard HJ. Cytolytic and antibacterial activity of synthetic peptides derived from amoebapore, the pore-forming peptide of Entamoeba histolytica. Proc Natl Acad Sci USA. 1994;91: 2602–2606.

17. Leippe M, Andrä J, Nickel R, Tannich E, Müller-Eberhard HJ. Amoebapores, a family of membranolytic peptides from cytoplasmic granules of Entamoeba histolytica: isolation, primary structure, and pore formation in bacterial cytoplasmic membranes. Mol Microbiol. 1994;14: 895–904.

18. Ravdin JI, Moreau F, Sullivan JA, Petri WA, Mandell GL. Relationship of free intracellular calcium to the cytolytic activity of Entamoeba histolytica. Infect Immun. 1988;56: 1505–1512.

19. Ralston KS, Solga MD, Mackey-Lawrence NM, Somlata, Bhattacharya A, Petri Jr WA. Trogocytosis by *Entamoeba histolytica* contributes to cell killing and tissue invasion. Nature. 2014;508: 526–530. doi:10.1038/nature13242

20. Ralston KS. Taking a bite: Amoebic trogocytosis in Entamoeba histolytica and beyond. Current Opinion in Microbiology. 2015;28: 26–35. doi:10.1016/j.mib.2015.07.009

21. Brown T. Observations by immunofluorescence microscopy and electron microscopy on the cytopathogenicity of naegleria fowleri in mouse embryo-cell cultures. Journal of Medical Microbiology. 1979;12: 363–371. doi:10.1099/00222615-12-3-363

22. Waddell DR, Vogel G. Phagocytic behavior of the predatory slime mold, Dictyostelium caveatum. Cell nibbling. Exp Cell Res. 1985;159: 323–334.

23. Batista FD, Iber D, Neuberger MS. B cells acquire antigen from target cells after synapse formation. Nature. 2001;411: 489–494. doi:10.1038/35078099

24. Wakim LM, Bevan MJ. Cross-dressed dendritic cells drive memory CD8+ T-cell activation after viral infection. Nature. 2011;471: 629–632. doi:10.1038/nature09863

25. Davis CO, Kim K-Y, Bushong EA, Mills EA, Boassa D, Shih T, et al. Transcellular degradation of axonal mitochondria. Proc Natl Acad Sci USA. 2014;111: 9633–9638. doi:10.1073/pnas.1404651111

26. Weinhard L, di Bartolomei G, Bolasco G, Machado P, Schieber NL, Neniskyte U, et al. Microglia remodel synapses by presynaptic trogocytosis and spine head filopodia induction. Nat Commun. 2018;9: 1228. doi:10.1038/s41467-018-03566-5

27. Abdu Y, Maniscalco C, Heddleston JM, Chew T-L, Nance J. Developmentally programmed germ cell remodelling by endodermal cell cannibalism. Nat Cell Biol. 2016;18: 1302–1310. doi:10.1038/ncb3439

28. Steele S, Radlinski L, Taft-Benz S, Brunton J, Kawula TH. Trogocytosis-associated cell to cell spread of intracellular bacterial pathogens. Monack D, editor. eLife. 2016;5: e10625. doi:10.7554/eLife.10625

29. Mercer F, Ng SH, Brown TM, Boatman G, Johnson PJ. Neutrophils kill the parasite Trichomonas vaginalis using trogocytosis. PLoS Biol. 2018;16: e2003885. doi:10.1371/journal.pbio.2003885

30. Matlung HL, Babes L, Zhao XW, van Houdt M, Treffers LW, van Rees DJ, et al. Neutrophils Kill Antibody-Opsonized Cancer Cells by Trogoptosis. Cell Rep. 2018;23: 3946–3959.e6. doi:10.1016/j.celrep.2018.05.082

31. Velmurugan R, Challa DK, Ram S, Ober RJ, Ward ES. Macrophage-Mediated Trogocytosis Leads to Death of Antibody-Opsonized Tumor Cells. Mol Cancer Ther. 2016;15: 1879–1889. doi:10.1158/1535-7163.MCT-15-0335

32. Miyake K, Shiozawa N, Nagao T, Yoshikawa S, Yamanishi Y, Karasuyama H. Trogocytosis of peptide-MHC class II complexes from dendritic cells confers antigen-presenting ability on basophils. Proc Natl Acad Sci USA. 2017;114: 1111–1116. doi:10.1073/pnas.1615973114

33. Gu P, Gao JF, D’Souza CA, Kowalczyk A, Chou K-Y, Zhang L. Trogocytosis of CD80 and CD86 by induced regulatory T cells. Cell Mol Immunol. 2012;9: 136–146. doi:10.1038/cmi.2011.62

34. Rossi EA, Goldenberg DM, Michel R, Rossi DL, Wallace DJ, Chang C-H. Trogocytosis of multiple B-cell surface markers by CD22 targeting with epratuzumab. Blood. 2013;122: 3020–3029. doi:10.1182/blood-2012-12-473744

35. Martínez-Martín N, Fernández-Arenas E, Cemerski S, Delgado P, Turner M, Heuser J, et al. T cell receptor internalization from the immunological synapse is mediated by TC21 and RhoG GTPase-dependent phagocytosis. Immunity. 2011;35: 208–222. doi:10.1016/j.immuni.2011.06.003

36. Gutiérrez-Kobeh L, Cabrera N, Pérez-Montfort R. A Mechanism of Acquired Resistance to Complement-Mediated Lysis by Entamoeba histolytica. The Journal of Parasitology. 1997;83: 234–241. doi:10.2307/3284446

37. Hamelmann C, Foerster B, Burchard GD, Shetty N, Horstmann RD. Induction of complement resistance in cloned pathogenic Entamoeba histolytica. Parasite Immunol. 1993;15: 223–228.

38. Hamelmann C, Urban B, Foerster B, Horstmann RD. Complement resistance of pathogenic Entamoeba histolytica mediated by trypsin-sensitive surface component(s). Infect Immun. 1993;61: 1636–1640.

39. Rastew E, Morf L, Singh U. Entamoeba histolytica rhomboid protease 1 has a role in migration and motility as validated by two independent genetic approaches. Exp Parasitol. 2015;154: 33–42. doi:10.1016/j.exppara.2015.04.004

40. Baxt LA, Rastew E, Bracha R, Mirelman D, Singh U. Downregulation of an Entamoeba histolytica Rhomboid Protease Reveals Roles in Regulating Parasite Adhesion and Phagocytosis. Eukaryot Cell. 2010;9: 1283–1293. doi:10.1128/EC.00015-10

41. Rigothier M-C, Khun H, Tavares P, Cardona A, Huerre M, Guillén N. Fate of Entamoeba histolytica during establishment of amoebic liver abscess analyzed by quantitative radioimaging and histology. Infect Immun. 2002;70: 3208–3215.

42. González-Ruiz A, Haque R, Aguirre A, Castañón G, Hall A, Guhl F, et al. Value of microscopy in the diagnosis of dysentery associated with invasive Entamoeba histolytica. J Clin Pathol. 1994;47: 236–239.

43. Capin R, Capin NR, Carmona M, Ortíz-Ortíz L. Effect of complement depletion on the induction of amebic liver abscess in the hamster. Arch Invest Med (Mex). 1980;11: 173–180.

44. Snow M, Chen M, Guo J, Atkinson J, Stanley SL. Differences in complement-mediated killing of Entamoeba histolytica between men and women--an explanation for the increased susceptibility of men to invasive amebiasis? Am J Trop Med Hyg. 2008;78: 922–923.

45. Costa CA, Nunes AC, Ferreira AJ, Gomes MA, Caliari MV. Entamoeba histolytica and E. dispar trophozoites in the liver of hamsters: in vivo binding of antibodies and complement. Parasit Vectors. 2010;3: 23. doi:10.1186/1756-3305-3-23

46. Reed SL, Curd JG, Gigli I, Gillin FD, Braude AI. Activation of complement by pathogenic and nonpathogenic Entamoeba histolytica. J Immunol. 1986;136: 2265–2270.

47. Lejeune A, Gicquaud C. Evidence for two mechanisms of human erythrocyte endocytosis by Entamoeba histolytica-like amoebae (Laredo strain). Biol Cell. 1987;59: 239–245.

48. Braga LL, Ninomiya H, McCoy JJ, Eacker S, Wiedmer T, Pham C, et al. Inhibition of the complement membrane attack complex by the galactose-specific adhesion of Entamoeba histolytica. J Clin Invest. 1992;90: 1131–1137. doi:10.1172/JCI115931

49. Ventura-Juárez J, Campos-Rodríguez R, Jarillo-Luna RA, Muñoz-Fernández L, Escario-G-Trevijano JA, Pérez-Serrano J, et al. Trophozoites of Entamoeba histolytica express a CD59-like molecule in human colon. Parasitol Res. 2009;104: 821–826. doi:10.1007/s00436-008-1262-3

50. Reed SL, Ember JA, Herdman DS, DiScipio RG, Hugli TE, Gigli I. The extracellular neutral cysteine proteinase of Entamoeba histolytica degrades anaphylatoxins C3a and C5a. J Immunol. 1995;155: 266–274.

51. Reed SL, Gigli I. Lysis of complement-sensitive Entamoeba histolytica by activated terminal complement components. Initiation of complement activation by an extracellular neutral cysteine proteinase. J Clin Invest. 1990;86: 1815–1822. doi:10.1172/JCI114911

52. Reed SL, Keene WE, McKerrow JH, Gigli I. Cleavage of C3 by a neutral cysteine proteinase of Entamoeba histolytica. J Immunol. 1989;143: 189–195.

53. Calderon J, Tovar R. Loss of susceptibility to complement lysis in Entamoeba histolytica HM1 by treatment with human serum. Immunology. 1986;58: 467–471.

54. Olivos-García A, Nequiz M, Liceaga S, Mendoza E, Zúñiga P, Cortes A, et al. Complement is a rat natural resistance factor to amoebic liver infection. Biosci Rep. 2018;38. doi:10.1042/BSR20180713

55. Huston CD, Boettner DR, Miller-Sims V, Petri, Jr. WA. Apoptotic Killing and Phagocytosis of Host Cells by the Parasite Entamoeba histolytica. Infect Immun. 2003;71: 964–972. doi:10.1128/IAI.71.2.964-972.2003

56. Sateriale A, Huston CD. A Sequential Model of Host Cell Killing and Phagocytosis by Entamoeba histolytica. J Parasitol Res. 2011;2011. doi:10.1155/2011/926706

57. Khalil MI, Foda BM, Suresh S, Singh U. Technical advances in trigger-induced RNA interference gene silencing in the parasite Entamoeba histolytica. Int J Parasitol. 2016;46: 205–212. doi:10.1016/j.ijpara.2015.11.004

